# Practical Use of Methods for Imputation of HLA Alleles from SNP Genotype Data

**DOI:** 10.1101/091009

**Authors:** Allan Motyer, Damjan Vukcevic, Alexander Dilthey, Peter Donnelly, Gil McVean, Stephen Leslie

## Abstract

The human leukocyte antigen (HLA) genes play an essential role in immune function. Typing of HLA alleles is critical for transplantation and is informative for many disease associations. The high cost of accurate lab-based HLA typing has precluded its use in large-scale disease-association studies. The development of statistical methods to type alleles using linkage disequilibrium with nearby SNPs, called HLA imputation, has allowed large cohorts of individuals to be typed accurately, so that massive numbers of affected individuals and controls may be studied. This has resulted in many important findings. Several HLA imputation methods have been widely used, however their relative performance has not been adequately addressed. We have conducted a comprehensive study to evaluate the most widely used HLA imputation methods. We assembled a multi-ethnic panel of 10,561 individuals with SNP genotype data and lab-based typing of alleles at 11 HLA genes at two-field resolution, and used it to train and validate each method. Use of this panel leads to imputation accuracy far superior to what is currently publicly available. We present a highly-accurate new imputation method, HLA*IMP:03. We address the question of optimal use of HLA imputations in tests of genetic association, showing that it is usually not necessary to apply a probability threshold to achieve maximal power. We also investigated the effect on accuracy of SNP density and population stratification at the continental level and show that neither of these are a significant concern.

## Introduction

Human leukocyte antigen (HLA) genes are immune-system genes that are of major biological and clinical interest. The genes are located in the major histocompatibility complex (MHC) on chromosome 6. HLA alleles are determinants of transplant compatibility,^1^ and have been associated with many conditions including autoimmune diseases (e.g. multiple sclerosis,^2,3^ ankylosing spondylitis,^4^ psoriasis,^5^ rheumatoid arthritis^6^), communicable diseases (e.g. cerebral malaria,^7^ HIV,^8^ enteric fever^9^), cancer (e.g. Hodgkin lymphoma,^10^ chronic lymphocytic leukemia,^11^) and adverse drug reactions.^12^ The alleles of these genes are expensive to type at high resolution using lab-based methods, meaning they are often neglected in studies of genetic association to disease and other phenotypes. A major advance was the development of statistical methods that use the correlation structure between HLA genes and nearby single nucleotide polymorphisms (SNPs) to type unknown HLA alleles from SNP array data.^13^ This process is known as HLA imputation. Methods that perform HLA imputation—which are high-throughput, accurate and relatively low cost—have enabled large-scale studies of genetic variation in the MHC, and significantly advanced the understanding of several diseases, e.g. Wellcome Trust Case Control Consortium 2 studies of multiple sclerosis,^2^ ankylosing spondylitis^4^ and psoriasis,^5^ and other studies.^3,6,14^

The first HLA imputation method, HLA*IMP^13,15^, has been used in many association studies.^2,4,5,10,16^ Subsequently other approaches have been developed, including HLA*IMP:02,^17^ HIBAG,^18^ SNP2HLA,^19^ MAGprediction^20^ and others.^21–24^ These methods have been applied to large cohort genetic studies.^2–6,14,25–30^ Much attention has recently turned to the typing of HLA alleles with NGS data,^31,32^ however HLA imputation with SNP genotypes remains an important and powerful tool for the study of the MHC, primarily because SNP genotyped datasets continue to have larger sample sizes (see e.g. Ref.^33^). The practical use of HLA imputation, including the relative performance of the available methods, has not been thoroughly addressed in the literature. To fully understand the utility and applicability of recent advances, we have undertaken a large study to explore the factors that affect the accuracy of HLA imputation and a comparison of four of the most widely used HLA imputation methods: HIBAG, HLA*IMP:02, SNP2HLA and MAGprediction. We did not evaluate HLA*IMP, as we consider it to have been superseded by HLA*IMP:02. We have also assessed the performance of the recently-developed method for imputation of KIR gene variation from SNP genotype data, KIR*IMP,^34^ when adapted to HLA genes, which we have named HLA*IMP:03 and made available as a web server (see Web Resources).

Each of the HLA imputation methods makes use of a sample (known as a “reference panel”) of individuals with known HLA and SNP genotypes. The methods involve a training step, in which a statistical model, relating HLA alleles to patterns of SNPs, is fitted to the reference panel; and an inference step, where HLA genotypes are imputed (predicted) using this model for a sample (known as a “study panel”) of individuals with known SNP genotypes but unknown HLA alleles. In practice, when testing or validating a method, the HLA alleles of the study panel will be known, but treated as missing, so that assessments of accuracy can be made. The accuracy of the imputations depends both on the data (the number of reference individuals and the genetic and ethnic diversity represented, the number of SNP loci used, how well the reference and study panels match, and the data quality) and the statistical method. The training step is typically computationally intensive and ordinarily only needs to be performed once for a given reference panel and HLA gene. Furthermore, access to reference panel data is often restricted. For these reasons HLA imputation methods are usually made available as pre-trained models that can be used with minimal effort to perform HLA imputation for a study panel without the need for training. In some circumstances, however, HLA imputation methods will need to be trained with a specific reference panel (e.g. if the study panel population is not well represented in the reference panel used for pre-training and the user has specific data they wish to use for the reference panel).

The methods we investigate, HIBAG, HLA*IMP:02, MAGprediction and SNP2HLA, have been previously compared with HLA*IMP (the only widely used method at the time these methods were published) using various reference and study panels,^15,18,20,26^ but have not been directly compared with each other. For most applications of HLA imputation it is the performance of the available pre-trained methods that is of relevance, however direct comparisons of the pre-trained methods (e.g. Ref.^35^) have been limited due to the difficulty in obtaining independent validation datasets that have not been used to train at least one of the methods. Further, the aggregation of individuals into larger reference panels, and especially the inclusion of more non-European individuals, means that previously published estimates of imputation accuracy do not reflect the performance currently available. We have assessed the performance of currently available pre-trained HLA imputation methods using an independent European validation dataset, providing a guide to current best practice.

The assessment of pre-trained HLA imputation methods is of practical relevance, but it does not allow the differences in performance to be attributed to the different statistical methods (since methods have been trained with different reference panels). To allow this comparison we evaluated the HLA imputation methods by training each with the same reference panel. Moreover, we have assembled the largest HLA reference panel to-date to perform this evaluation, meaning that the findings establish the best general imputation performance currently possible among the leading methods.

By combining several previously collected sample sets (see Material and Methods) we assembled a multi-ethnic panel of 10,561 individuals (8,768 European, 869 Asian, 568 African-American/African and 356 Latino) with SNP genotype data and lab-based typing of alleles at 11 HLA genes at two-field resolution (which specifies a unique amino acid sequence; formerly referred to as ‘four-digit’ resolution). Note that not all samples are typed at all HLA loci (see Material and Methods). The performance of each method was evaluated via five-fold cross-validation (CV) using the multi-ethnic dataset, and also using only European individuals to investigate the effect of population stratification at the continental level. To assess the effect of potential differences in the collection and typing protocols for the constituent sub-sets of the main data we also carried out an analysis using European individuals partitioned into independently collected and genotyped reference and validation sets. We assessed both the accuracy and calibration of each method. We also performed CV to assess the performance of the newly-presented method, HLA*IMP:03, which we found to be the most accurate, when the study panel is typed on a number of different commercial SNP arrays with differing SNP content.

Finally, we consider the optimal use of HLA imputations in downstream analyses. Each method provides a probability distribution over all possible genotypes at the imputed locus, which reflects the confidence in the imputation. The imputed genotype is the mode of this probability distribution, which we refer to as the *maximum a posteriori* (MAP) imputation. Subsequent analyses that make use of the MAP imputations can then be restricted to those with probability greater than a specified call threshold. Typically, the imposition of a call threshold will result in an improvement in accuracy for those imputations that are called, at the expense of a lower call rate. In assessing the performance of HLA imputation, previous studies have presented the accuracy and call rate at call thresholds that have been set without a rigorous analysis (at 0.7 for HLA*IMP and HLA*IMP:02, or 0.5 for HIBAG). We consider the effect of the call threshold on the power of statistical tests of association that use the MAP imputation. An alternative approach, that we also consider, is to use the expected genotype score or ‘dosage’ for each possible allele. We compare methods on the basis of the optimal power achieved by these approaches. This not only provides a more meaningful way of comparing methods, but provides a guide to the optimal use of HLA imputations. In the association study context, we show that the use of dosages usually attains similar power to MAP imputations, and if MAP imputations are used then it is usually not necessary to apply a call threshold to attain optimal power.

## Material and Methods

### DNA Samples and HLA Typing

We assembled a large dataset by merging data from a number of sources. The full dataset consisted of 10,561 individuals. Categorized by self-reported ancestry there were 568 African-American/African, 869 Asian, 8,768 European, and 356 Latino individuals. Each individual was typed at two-field resolution at both alleles for at least one of 11 HLA loci (listed in Table 1) and at SNPs in the extended MHC (xMHC; defined as the region on chromosome 6 spanning 26.033.5 Mb according to GRCh37 coordinates). The merged dataset was assembled from the following sources: the 1958 Birth Cohort (see Web Resources), HapMap CEU individuals^36^ and CEPH CEU+ additional individuals^37^ with HLA typing as described in Ref.^17^ and quality control (QC) procedures applied as described in Ref.^15^ (referred to collectively in this paper as “CEU+58”) a Glaxo-SmithKline dataset (here labeled “GSK”, also known as HLARES in other publications) and HapMap YRI individuals^36^ (“YRI”, as described in Ref.^17^ and QC as described in Ref.;^15^ 1000 Genomes Project^38,39^ (“1000”) with standard HLA*IMP dataset QC with the HLA*IMP interface, using a 20% missing data threshold on SNPs and individuals (see Web Resources); individuals obtained from the Type 1 Diabetes Genetics Consortium^19,40^ (“T1DGC” as used in Ref.;^19^ an unpublished dataset of African-Americans provided by colleagues at Kings College, London (“KC”), with Sequence Based Typing (SBT) of HLA performed at Oklahoma Medical Research Foundation and University of Alabama and SNP QC as for the 1000G dataset with a missing data threshold of 5%; a pan-Asian dataset (“PA”) comprising three Southeast Asian populations sampled from Singapore and HapMap (CHB and JPT) individuals^27,41^ (see Web Resources); and an unpublished dataset of Swedish individuals provided by Karolinska Institutet (“SW”) with high-resolution HLA typing (SBT) and QC as for the KC dataset.

**Table 1.** Summary of HLA reference panel. For each HLA locus the table displays: the number of individuals typed at two-fields in each dataset (CEU+58, GSK, YRI, 1000G, T1DGC, KC, PA, SW) and in total; the number of SNPs in the xMHC for the dataset formed by merging all individuals typed at that HLA locus; and the number of two-field alleles present in the dataset. MIM number – Mendelian Inheritance in Man number. *HLA-DRB4* does not have a MIM number; the HUGO Gene Nomenclature Committee (HGNC) ID is given instead.

| HLA locus | *A* | *B* | *C* | *DQA1* | *DQB1* | *DRB1* | *DRB3* | *DRB4* | *DRB5* | *DPA1* | *DPB1* |
| --- | --- | --- | --- | --- | --- | --- | --- | --- | --- | --- | --- |
| MIM number or HGNC ID | 142800 | 142830 | 142840 | 146880 | 604305 | 142857 | 612735 | HGNC:4952 | 604776 | 142880 | 142858 |
| *No. individuals in each dataset* |  |  |  |  |  |  |  |  |  |  |  |
| CEU+58 | 890 | 1,490 | 836 | 61 | 1,017 | 1,122 | – | – | – | – | – |
| GSK | 379 | 1,365 | 398 | 302 | 498 | 1,191 | 158 | 182 | 265 | – | 112 |
| YRI | 23 | 23 | 23 | 23 | 22 | 18 | – | – | – | – | – |
| 1000G | 927 | 927 | 926 | – | 925 | 927 | – | – | – | – | – |
| T1DGC | 5,192 | 5,192 | 5,192 | 5,192 | 5,192 | 5,192 | – | – | – | 5,192 | 5,190 |
| KC | – | – | – | 332 | 330 | 331 | 332 | 332 | 332 | – | – |
| PA | 443 | 439 | 449 | 450 | 450 | 448 | – | – | – | 452 | 441 |
| SW | – | – | – | 423 | 421 | 423 | 48 | 42 | 9 | 423 | 422 |
| Total individuals | 7,854 | 9,436 | 7,824 | 6,783 | 8,855 | 9,652 | 538 | 556 | 606 | 6,067 | 6,165 |
| No. SNPs in merged dataset | 1,672 | 1,672 | 1,672 | 1,357 | 1,162 | 1,162 | 2,254 | 2,253 | 2,291 | 2,193 | 2,097 |
| No. two-field alleles | 91 | 174 | 54 | 20 | 28 | 94 | 6 | 3 | 4 | 8 | 51 |

The constituent datasets were typed on various SNP arrays, as listed in Table S1. In addition to the QC applied individually to the data from each source, we performed further QC steps to allow us to combine these datasets into a single reference panel. This included: converting the SNPs to GRCh37 coordinates using liftOver (see Web Resources); converting to a common strand alignment using PLINK (see Web Resources); and excluding SNPs that could not be aligned or had very different allele frequencies between the datasets. As part of the merging process we excluded duplicate individuals (there was overlap between some of the datasets, e.g. CEU and 1000G). We treated ambiguous HLA types, e.g. G and P coded alleles (see Web Resources), as the most frequent allele, and did not attempt to account for differences in HLA allele nomenclature between the constituent datasets. The number of individuals and the number of SNPs in the xMHC passing QC for each data source is shown in Table S1. We adopted two different approaches to the merging of SNP data. Firstly, we merged datasets using overlapping SNP genotypes. Secondly, we merged datasets after imputing SNPs within each dataset. In the second merged dataset with imputed SNPs, haplotypes were phased (required by HLA*IMP:03). We produced two sets of phasings: SNPs and HLA alleles phased together (used as a reference panel) and SNPs phased without HLA (used for validation). The procedure for merging datasets is shown in Figure S1 and described below.

The merged dataset with overlapping SNPs was created separately for each HLA locus because of the differing extent of lab-based typing. Specifically, for each locus we included only the individuals which had lab-based HLA types for that locus, and only the SNPs that were polymorphic and were typed in at least 98% of that set of individuals. The number of individuals and number of SNPs passing QC in the merged dataset with overlapping SNPs at each HLA locus is shown in Table 1 and Table S2.

The merged dataset with imputed SNPs was formed by first imputing SNPs within each dataset with the Michigan Imputation Server (imputation with Minimac3,^42^ see Web Resources), selecting the appropriate population for QC, SHAPEIT^43^ for pre-phasing, and 1000 Genomes Phase 3 version 5 as the reference panel. Individuals in each dataset belonging to each of the 1000 Genomes “super populations” (Ad Mixed American, African, East Asian, European) were imputed separately, to make full use of the available population-specific QC, and use all genotyped SNPs (not just overlapping SNPs). The datasets with separately imputed SNP genotypes were merged by retaining only biallelic SNPs directly genotyped in at least one dataset and with R-squared output by Minimac3 greater than 0.8 in all datasets. We used a cut-off more stringent than a standard cut-off of 0.3 as inspection of the distribution of R-squared values showed that this had a small effect on the number of SNPs retained. We did not filter imputed SNPs on MAF < 0.01 as usual as we were combining samples of different ethnicity and had already restricted the imputed SNPs to those directly genotyped in at least one dataset. We then retained SNPs with at least 95% of posteriors greater than 0.9 in all imputed datasets, and at least 98% of posteriors greater than 0.9 in the merged dataset. We inspected the distribution of posteriors to set the above stringent cut-offs. This resulted in a single merged dataset to be used for all HLA loci with 6,452 SNPs across the xMHC.

The merged dataset with imputed SNPs was phased (using only SNP genotypes, i.e. without HLA types) using SHAPEIT (version 2) and the 1000 Genomes Phase 3 reference panel. These phased SNPs were to be used for validation, so 916 samples that were in both the merged dataset and the phasing reference panel were removed from the reference panel, to mirror the typical situation where study samples do not have known phase.

We, finally, phased haplotypes of HLA alleles and SNPs in the merged dataset with imputed SNPs. This was performed separately for each HLA locus due to the different extent of HLA typing for each locus. There were some samples with previously determined phasing of HLA alleles and SNPs (87 CEU+58 and 60 YRI individuals, see Ref.^13^). We attempted to resolve the phase of HLA alleles and the imputed SNPs for these samples. To do this, the imputed SNPs were phased with SHAPEIT using the 1000 Genomes Phase 3 reference panel, or in the case of samples appearing in the reference panel we used the reference haplotypes. We then matched these phased SNP haplotypes (with imputed SNPs) with the original SNP haplotypes (with matching HLA phase) by inspection of the number of identical SNPs. This resolved the phasing of HLA alleles in the haplotypes with imputed SNPs in 67 CEU+58 and 56 YRI individuals, with those not resolved likely due to phasing switch errors. We then attempted to resolve the phase of HLA alleles and SNPs in all remaining samples. SHAPEIT was used with the 123 samples with resolved HLA phase as a reference panel. Each HLA locus was phased separately using only samples with HLA typed at both alleles. Due to the limitations of SHAPEIT, each allele of an HLA gene was represented as a separate genetic variant, coded on the basis of presence/absence, and with position at the center of the HLA locus. SHAPEIT was run with the number of conditioning states set to 1000 (increased from default of 100). Since each HLA allele was represented as a separate variant it was possible for the two HLA alleles of an individual to be erroneously phased to the same haplotype. Across the 11 HLA loci, this occurred in at most 0.3% of samples, and any individual for which this occurred was discarded from the dataset.

### HLA Imputation

HIBAG version 1.4.0 (with R statistical software version 3.2.0) was used with SNPs in the recommended flanking region 500 kb each side of the HLA locus, and all default settings (including 100 classifiers). HIBAG was not run for *HLA-DRB3* and -*DRB4* as it does not provide functionality to select the nearby SNPs to be used for imputation at these HLA loci.

HLA*IMP:02 was run with parameter settings modified from those described in its original publication^17^ to accommodate the large number of individuals in the reference panel, which has more than doubled since the previous study, while achieving feasible computational and memory requirements (a few days of running time for training on a high-performance computing cluster and less than 250 GB memory). Our investigations using moderately sized reference samples showed that the parameters could be modified to run faster and use less memory without noticeable loss of accuracy and that any potentially small decrease in accuracy due to changes in the settings is compensated by the ability to handle a larger reference panel (data not shown). The modified settings (refer to the original paper for definitions^17^) were: localization feature turned off (previously turned on at all loci except *HLA-B* and-*DRB1*); graph sampling error, *m_S_*, and graph building error, *m_B_*, probabilities both set to 0.001 (previously both were set to 0.002); and the number of sampled haplotype pairs, *N_S_*, set to 5 (previously 50). We also made changes to how the SNPs used in imputation were selected. Previously, HLA*IMP:02 made use of the nearest 300 SNPs on either side of the center of the HLA locus, excluding any SNPs that gave a p-value less than 10^−5^ on a test for deviation from Hardy-Weinberg equilibrium (HWE). Here we included the nearest 300 SNPs on either side of the center of the HLA locus and additionally all SNPs in the region flanking 500 kb either side of the HLA locus, and did not exclude any SNPs due to deviation from HWE. The inclusion of additional SNPs in the region flanking 500 kb either side of the HLA locus was made to allow a sufficiently wide SNP window in regions of high SNP density. It resulted in the inclusion of all SNPs used by HIBAG, but this was only an additional 12 SNPs at *HLA-B*, 53 at *HLA-DPA1*, 35 at *HLA-DPB1*, and none at the other HLA loci. We also ran a version of HLA*IMP:02 with the reference panel set as pre-phased haplotypes, and all other settings unchanged (referred to below as ‘HLA*IMP:02 (phased)’).

HLA*IMP:03 was applied to each HLA locus using phased haplotypes with SNPs within 500 kb either side of each locus, 400 trees, and with the parameter *m* (the number of SNPs randomly sampled as candidates for each split of each tree) set to 100. For training, we used the version of the data where both SNPs and HLA alleles were phased. For inference, we used the version of the data where only the SNPs were phased (to more properly mimic how real study data would be prepared).

The command line version of MAGprediction was run (with Matlab Compiler Runtime version 7.14).

SNP2HLA version 1.0.3 (with PLINK version 1.07,^44^ and Beagle version 3.0.4^45^) was used. The ‘MakeReference’ script was run to perform training, followed by the ‘SNP2HLA’ script for inference, both with window size set to the default of 1000.

### Assessing Imputation Performance

We conducted a number of experiments, described below, to assess the performance of the HLA imputation methods. In each experiment imputed HLA alleles are compared with the known, lab-derived, HLA types at two-field resolution, with the exception of *HLA-DQA1*. At this locus we assessed performance at the resolution of G groups. That is, imputations were considered correct if they belonged to the same G allele group as the lab-derived HLA type. This was done because the T1DGC samples were typed at the resolution of G groups, but the other samples were typed at higher resolution without ambiguous G alleles. (At this locus 12.0% of alleles in the non-T1DGC samples were not present in the T1DGC samples, instead represented as the most common allele in the G group.) There wasn’t evidence of this issue at other HLA loci.

Each method reports a posterior probability for imputed HLA alleles. These probabilities are potentially reflective of the confidence one can ascribe to a given imputation (see the assessment of calibration below). HLA*IMP:02, HLA*IMP:03 and SNP2HLA produce a posterior probability for each imputed allele, i.e. two probabilities for each imputed genotype (although this is not provided directly by SNP2HLA), whereas HIBAG and MAGprediction produce a single probability for each imputed genotype as a whole. Although the probabilities are not directly comparable, to enable some comparison we treated the HIBAG and MAGprediction probabilities as applying individually to each allele. For a given call threshold, *T*, 0 ≤ *T* ≤ 1, the accuracy of imputations was calculated as the proportion of those imputed alleles with posterior greater than or equal to *T* that match the known HLA types. In the case of an imputed homozygous genotype, only the allele with the greater posterior was considered correct if only one copy of the imputed allele is present in the lab-based genotype (i.e. where the latter is heterozygous). The associated call rate was calculated as the proportion of imputed alleles with posterior probability greater than or equal to *T*. For each estimate of accuracy, we calculated a 95% Bayesian credible interval on the basis of a binomial model and a uniform prior distribution (in the context of cross-validation this should be treated as an approximation only, see Ref.^34^ for further discussion).

We measure the impact of using HLA imputations on the power of a statistical test of association, where each allele is tested by coding its absence and presence as 0 and 1, respectively, and the genotype of each individual is coded as 0, 1 or 2 (i.e. a single allele is compared against all other alleles). The situation is then analogous to that for a biallelic marker. The most useful measure of the effectiveness of imputation in this scenario is the *effective sample size*. The effective sample size with perfectly typed alleles has power equivalent to that of the actual study sample size with (imperfect) imputations. A more convenient quantity, which we use, is the *sample size ratio*, the ratio of the effective sample size to the actual study sample size. Under certain assumptions (see Appendix A) the sample size ratio is given by the square of the Pearson correlation coefficient between the true genotypes and MAP imputed genotypes (*r*^2^). When a call threshold *T* is imposed it is apparent that the quantity *r*^2^ should be replaced by 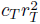, where *c_T_* is the call rate at threshold *T* and 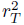 is based on only those imputations that are called (see Appendix A). An alternative to the use of the MAP imputed genotypes is the imputed dosage,^46,47^ the expected number of copies of each allele carried by each individual (a continuous value between 0 and 2). If MAP imputed genotypes are replaced by imputed dosages for each allele then the relevant quantity is the *r*^2^ between the true genotypes and imputed dosages. To assess the performance of the imputation methods in terms of power to detect associations for each allele we computed the sample size ratio for MAP imputations for a range of call thresholds and for imputed dosages.

A desirable property of imputation methods is that they be ‘well-calibrated’, in the sense that for all imputations with associated probabilities of, for example, 0.9, 90% of such imputations are correct. The idea is that the probabilities reported by the model may be meaningfully interpreted. We assessed the calibration of the methods by binning the imputed alleles by their posterior probabilities and comparing the mean probability in each bin to the imputation accuracy for that bin.

### Validation experiments

A number of validation experiments were performed, as listed in Table S3 and now described. Each validation experiment was performed separately for each HLA locus. We first compared the accuracy of HLA imputation methods using an identical reference panel. We partitioned the dataset into a reference panel and a study panel (here referred to as a “validation set” as the HLA types for these samples are known, and are used for comparison to the imputed HLA types) multiple times, as described below, allowing for several replications of HLA imputation testing using different reference and validation sets. We assessed the accuracy of each HLA imputation method on each reference and validation set pair. Unless otherwise stated, both the reference and validation set were restricted to individuals with both alleles typed to two-field resolution at the HLA locus. Specifically, we performed a five-fold CV, whereby the individuals are divided randomly into five sets (called “folds”), training is performed using four of the folds as the reference set and the remaining fold is imputed and used for testing accuracy. This is repeated five times, each time with a different fold as the validation set, resulting in a single imputation being made for each sample.

Five-fold CV was performed for a number of different versions of the merged dataset. The first CV experiment used the merged dataset with overlapping SNPs, and included all individuals (i.e. we used a multi-population reference panel). Despite considerable effort, we were unable to train MAGprediction on the CV dataset, and have therefore excluded it from this and the subsequent CV experiments. The version of HLA*IMP:02 with phased reference panel and HLA*IMP:03 made use of the phased haplotype version of the merged dataset. The phasing was performed using imputed SNPs, however here the SNPs were restricted to those used for the other methods (i.e. imputed SNPs were excluded for a fair comparison). The validation sets for HLA*IMP:03 made use of the haplotype phasing that was performed without HLA types.

The dataset included a number of individuals that were typed to two-field resolution for only one HLA allele (see Table S2). In order to test the effect of their inclusion, for HLA*IMP:02 and SNP2HLA we repeated CV with these extra individuals added to the reference panel, but not to the validation set, and we omitted the second allele (i.e. treated it as missing data). When running SNP2HLA in this manner for *HLA-B*, we found that two of the five folds resulted in the required memory exceeding 500 GB; we reduced the window size to 500 for these two folds. This extra analysis was not run with HIBAG because it can only handle reference individuals that are typed at both alleles, and HLA*IMP:03 was not run as haplotype phasing was not available for these individuals.

We repeated CV using the same assignment of individuals to folds in the first CV above, but with non-European individuals excluded. By comparing to the results of the first analysis, we aimed to investigate the effect of the presence of non-European individuals in the reference panel on imputations for European individuals, and more generally to test the effect of using a multi-population reference panel on imputation accuracy for study panels derived from a single population.

The random partitioning of individuals for CV means that individuals from each data source are likely to be represented in both the training and validation sets. Thus, any biases, or indeed systematic errors, present in one of the constituent data sources are likely to be represented in both the reference and validation datasets, potentially somewhat ameliorating their effect on the validation study. We therefore carried out a validation analysis in which the training and validation sets were composed exclusively of different data sources (i.e. independently ascertained). This is closer to the situation encountered by users of HLA imputation in practice. Training was performed using the T1DGC dataset (5,192 European individuals) and inferences made on all other European individuals (3,587 individuals). This analysis was carried out with the eight HLA loci for which the T1DGC data is typed (see Table 1).

We next assessed the performance of pre-trained models (i.e. methods not necessarily trained with the same reference panel) by using the T1DGC dataset (composed of 5,192 European individuals, 7,135 SNPs typed on Illumina HumanImmuno BeadChip) as the validation set. For HIBAG we used the pre-trained ImmunoChip European model (“ImmunoChip-European-HLA4-hg18.RData”; n.b. separate models are available for different SNP arrays and populations), with assembly set to “hg18”(which was the assembly of both the model and validation set), SNP matching based on position, and other options set to their default values. Across the seven HLA loci imputed, between 4.1% and 16.8% of SNPs in the HIBAG model were missing from the validation set. Pre-trained MAGprediction was run using the “General” models (which use a general set of HapMap SNPs), following instructions to first impute SNPs in the validation set using IMPUTE2 (v2.3.2)^48^ with the provided genetic map and reference panel. SNP2HLA is not currently available with a European reference panel so we did not include it in this comparison. The currently available versions of HLA*IMP:02 (available via Affymetrix as Axiom HLA Analysis software, see Web Resources) and HLA*IMP:03 make use of the entire merged dataset (including T1DGC), so we cannot use them directly to assess performance in the T1DGC dataset. Instead, we assessed two sets of imputations: (i) the CV experiment using the multi-population reference panel described above, and (ii) imputation with all T1DGC samples removed from the reference panel.

We carried out additional analyses to investigate the performance of HLA*IMP:03 in practical settings, since it was found to be the most accurate in the above analyses and is newly presented in this paper. The effect of increased SNP density in the reference panel through the use of imputed SNPs was investigated. As HLA imputation reference panels increase in size with the addition of datasets possibly typed on different SNP arrays, the overlap between SNPs typed in reference individuals may decrease, potentially leading to decreased imputation accuracy. SNP imputation, however, can be used to merge datasets without a decrease in SNP density. We performed five-fold CV with HLA*IMP:03 using the merged dataset with imputed SNPs.

We also investigated the performance of HLA*IMP:03 when using only SNPs typed on common SNP arrays, which is of practical relevance as study panels will typically be typed on a specific array. We considered a number of different Illumina and Affymetrix arrays (listed in Table S6), obtaining a list of SNPs for each one from the manufacturers’ websites (see Web Resources). For each array, we trained HLA*IMP:03 using the merged dataset with imputed SNPs, but with SNPs restricted to those typed on the array by matching on the GRCh37 genomic coordinates, and used the out-of-bag (OOB) estimate of imputation accuracy (the latter was done for computational convenience; see Ref.^34^ for a description of OOB and a comparison to CV). This choice of SNP set mimics the (best-case) scenario where all the SNPs in the intersection are perfectly typed in the study panel; in practice, typing for some SNPs will most likely fail, leading to possibly lower imputation accuracy. For this analysis, when the number of SNPs in the intersection was fewer than 300, we set *m* to be one third of that number (but always at least *m* = 1).

## Results

### Cross-validation accuracy

The accuracy (calculated with call threshold *T* = 0, unless otherwise stated) of HLA imputation with each method at two-field resolution for five-fold CV using all individuals typed at two-fields at both alleles at each of 11 HLA loci (*HLA-A*, -*B*, -*C*, -*DQA1*, -*DQB1*, -*DRB1*, -*DRB3*, -*DRB4*, -*DRB5*, -*DPA1* and -*DPB1*) was calculated for individuals in each of the four major populations in the dataset, and is presented in Table 2 and Figure 1. Results for 13 populations in the 1000G dataset are included in Table S4. HIBAG, HLA*IMP:02 and HLA*IMP:03 had similar imputation accuracy (shown by overlapping credible intervals) across each of the HLA loci and populations with a few exceptions: HIBAG performed worse at *HLA-A* in Asians; HLA*IMP:02 performed worse at *HLA-B* in Asians and Europeans, and at *HLA-DQB1* in Europeans; and HLA*IMP:03 was superior at *HLA-DRB1*. Despite the similarity in performance of these methods, there tended to be an ordering from best to worse of HLA*IMP:03, HIBAG, HLA*IMP:02. This ordering is also consistent with the deviations from similar performance listed above. SNP2HLA consistently performed substantially worse than the other three methods. Imputations for Europeans were more accurate than for non-Europeans, which was expected given the greater number of European individuals in the reference panel.

**Figure 1.**
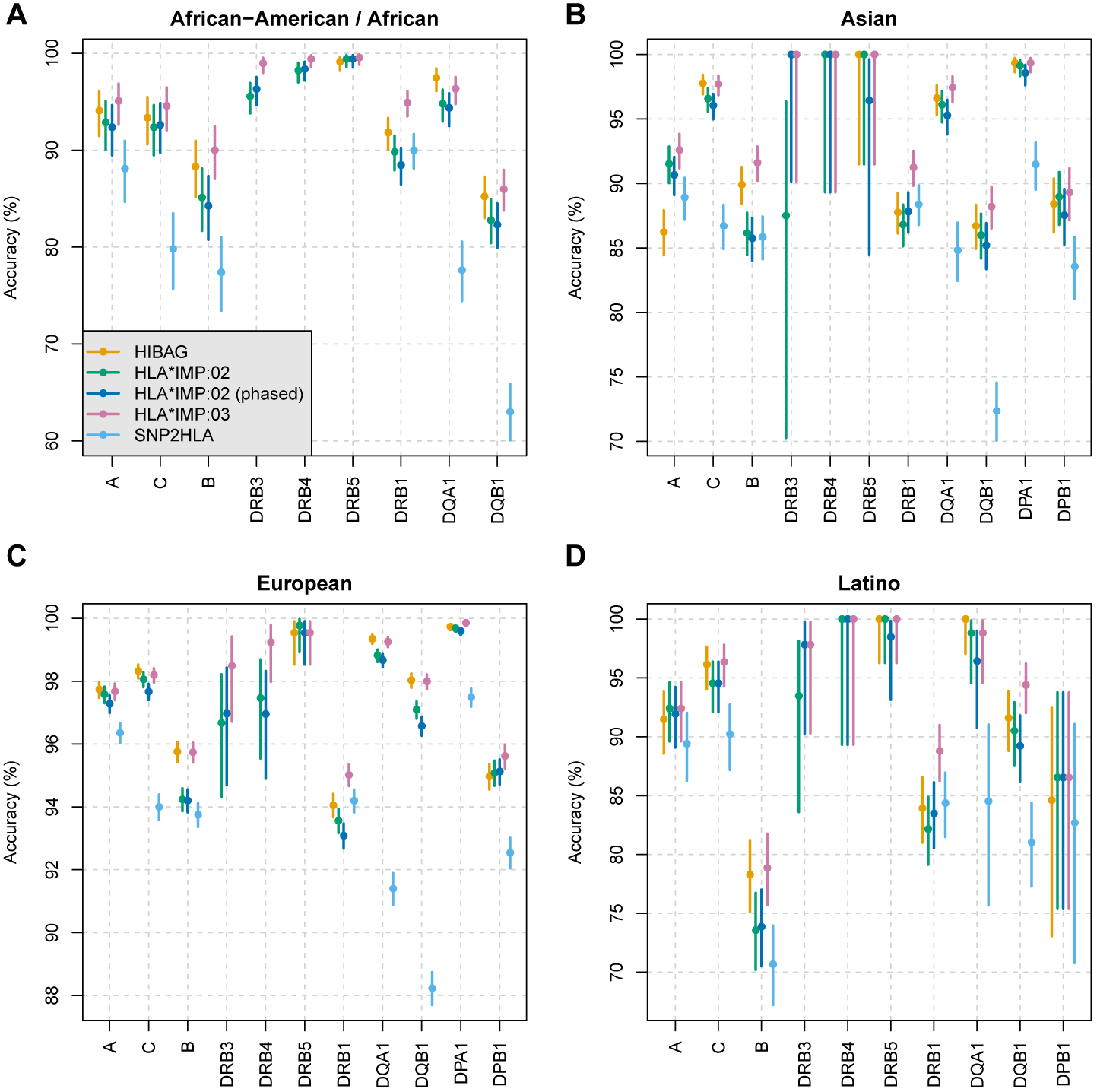
Cross-validation imputation accuracy. The percentage of correctly imputed two-field alleles for the different methods at each HLA locus from the cross-validation analysis with multi-population reference panel, with associated 95% credible intervals (see Material and Methods) for (**A**) African-American/African individuals, (**B**) Asian individuals, (**C**) European individuals, and (**D**) Latino individuals.

**Table 2.** Cross-validation imputation accuracy. The percentage of correctly imputed two-field alleles is shown for each method at each HLA locus in each of four major populations from the cross-validation analysis with the multi-population reference panel. The number of individuals used for validation in each population at each HLA locus is also shown. *Method does not support this locus.

| Population | HLA locus | No. validation individuals | Method |  |  |  |  |
| --- | --- | --- | --- | --- | --- | --- | --- |
|  |  |  | HIBAG | HLA*IMP:02 | HLA*IMP:02 (phased) | HLA*IMP:03 | SNP2HLA |
| African-American / African | *A* | 203 | 94.1 | 92.9 | 92.4 | 95.1 | 88.1 |
|  | *B* | 235 | 88.3 | 85.1 | 84.3 | 90.0 | 77.4 |
|  | *C* | 203 | 93.3 | 92.4 | 92.6 | 94.6 | 79.8 |
|  | *DQA1* | 355 | 97.5 | 94.8 | 94.4 | 96.3 | 77.6 |
|  | *DQB1* | 531 | 85.2 | 82.8 | 82.3 | 86.0 | 63.0 |
|  | *DRB1* | 550 | 91.8 | 89.8 | 88.5 | 94.9 | 90.0 |
|  | *DRB3* | 338 | * | 95.6 | 96.3 | 99.0 | * |
|  | *DRB4* | 337 | * | 98.2 | 98.4 | 99.4 | * |
|  | *DRB5* | 342 | 99.1 | 99.4 | 99.4 | 99.6 | * |
| Asian | *A* | 749 | 86.2 | 91.5 | 90.7 | 92.6 | 88.9 |
|  | *B* | 852 | 89.9 | 86.2 | 85.7 | 91.6 | 85.8 |
|  | *C* | 759 | 97.8 | 96.6 | 96.0 | 97.7 | 86.7 |
|  | *DPA1* | 452 | 99.3 | 99.1 | 98.6 | 99.3 | 91.5 |
|  | *DPB1* | 452 | 88.4 | 89.0 | 87.5 | 89.3 | 83.6 |
|  | *DQA1* | 487 | 96.6 | 96.1 | 95.3 | 97.4 | 84.8 |
|  | *DQB1* | 767 | 86.7 | 86.0 | 85.2 | 88.2 | 72.4 |
|  | *DRB1* | 845 | 87.8 | 86.8 | 87.8 | 91.2 | 88.4 |
|  | *DRB3* | 12 | * | 87.5 | 100.0 | 100.0 | * |
|  | *DRB4* | 11 | * | 100.0 | 100.0 | 100.0 | * |
|  | *DRB5* | 14 | 100.0 | 100.0 | 96.4 | 100.0 | * |
| Europeans | *A* | 6,685 | 97.7 | 97.6 | 97.3 | 97.7 | 96.4 |
|  | *B* | 7,999 | 95.8 | 94.2 | 94.2 | 95.7 | 93.7 |
|  | *C* | 6,642 | 98.3 | 98.1 | 97.7 | 98.2 | 94.0 |
|  | *DPA1* | 5,615 | 99.7 | 99.7 | 99.6 | 99.9 | 97.5 |
|  | *DPB1* | 5,686 | 95.0 | 95.1 | 95.1 | 95.6 | 92.5 |
|  | *DQA1* | 5,899 | 99.3 | 98.8 | 98.7 | 99.3 | 91.4 |
|  | *DQB1* | 7,325 | 98.0 | 97.1 | 96.6 | 98.0 | 88.2 |
|  | *DRB1* | 7,918 | 94.1 | 93.6 | 93.1 | 95.0 | 94.2 |
|  | *DRB3* | 165 | * | 96.7 | 97.0 | 98.5 | * |
|  | *DRB4* | 197 | * | 97.5 | 97.0 | 99.2 | * |
|  | *DRB5* | 217 | 99.5 | 99.8 | 99.5 | 99.5 | * |
| Latino | *A* | 217 | 91.5 | 92.4 | 91.9 | 92.4 | 89.4 |
|  | *B* | 350 | 78.3 | 73.6 | 73.9 | 78.9 | 70.7 |
|  | *C* | 220 | 96.1 | 94.5 | 94.5 | 96.4 | 90.2 |
|  | *DPB1* | 26 | 84.6 | 86.5 | 86.5 | 86.5 | 82.7 |
|  | *DQA1* | 42 | 100.0 | 98.8 | 96.4 | 98.8 | 84.5 |
|  | *DQB1* | 232 | 91.6 | 90.5 | 89.2 | 94.4 | 81.0 |
|  | *DRB1* | 339 | 83.9 | 82.2 | 83.5 | 88.8 | 84.4 |
|  | *DRB3* | 23 | * | 93.5 | 97.8 | 97.8 | * |
|  | *DRB4* | 11 | * | 100.0 | 100.0 | 100.0 | * |
|  | *DRB5* | 33 | 100.0 | 100.0 | 98.5 | 100.0 | * |

To assess the effect of pre-phasing the data on imputation accuracy we ran HLA*IMP:02 with a training set of phased haplotypes (HLA*IMP:02 (phased)) rather than genotypes (HLA*IMP:02). This also allows us to understand whether the difference in accuracy between HLA*IMP:02 and HLA*IMP:03 can be accounted for by pre-phasing. As shown in Table 2 and Figure 1, pre-phasing did not result in significant differences in imputation accuracy for HLA*IMP:02. HLA*IMP:02 with pre-phasing was not considered in subsequent analyses.

Five-fold CV repeated for HLA*IMP:02 and SNP2HLA with additional training individuals typed at two-fields at only one allele (with the other allele set to missing), and with the same validation individuals, showed no change in accuracy (data not shown).

### European-only reference panel

The accuracy for European individuals at two-fields for five-fold CV when training with all individuals was compared against the accuracy obtained using the same folds with non-Europeans removed, as shown in Figure S2. Removing non-European individuals from the training sets did not result in an improvement in accuracy for European individuals at any locus for any method. Rather, it had either a negligible or slightly detrimental effect, the latter particularly for HLA*IMP:02 at *HLA-DPA1* and for SNP2HLA at many of the genes.

### Validation with independently ascertained samples

The accuracy for European individuals in the CEU+58, GSK, 1000G and SW datasets (the datasets other than T1DGC containing European individuals) after training each method on the T1DGC dataset is shown in Figure S3 and Table S5. Accuracy aggregated over all validation individuals is given in Figure S4. The results were consistent with those for Europeans in five-fold CV (Figure 1C).

### Pre-trained methods

The above results involve comparisons of the imputation methods when used with identical training data. Accuracy for the pre-trained methods, which each use different training data, with validation in T1DGC dataset, are presented in Table 3 and Figure 2. As mentioned in the Material and Methods section, SNP2HLA was not included in this analysis as it is not currently publicly available with a European reference panel.

**Figure 2.**
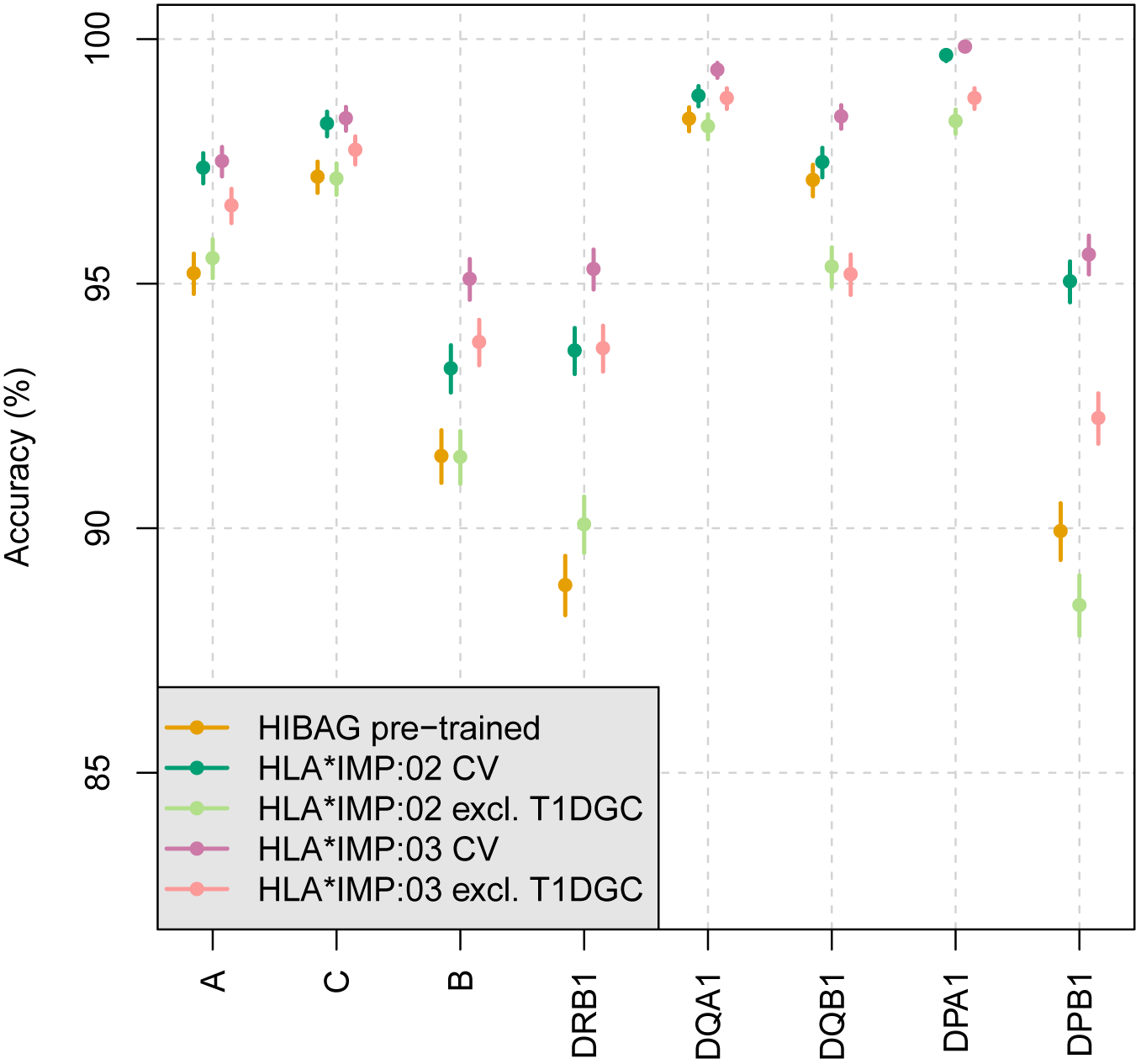
Imputation accuracy in T1DGC validation set. The percentage of correctly imputed two-field alleles for the different methods at each HLA locus in the T1DGC validation set, with associated 95% credible intervals (see Material and Methods). The publicly available pre-trained model was used for HIBAG. Two sets of results are presented for HLA*IMP:02 and HLA*IMP:03: the cross-validation analysis with the multi-population reference panel, and trained with all samples except T1DGC. Pre-trained MAGprediction was omitted from the figure due to its very low accuracy.

**Table 3.** Imputation accuracy in T1DGC validation set. The percentage of correctly imputed two-field alleles is shown for each method at each HLA locus in the T1DGC validation set. Publicly available pre-trained models were used for HIBAG and MAGprediction. Two sets of results are presented for HLA*IMP:02 and HLA*IMP:03: the cross-validation analysis with the multi-population reference panel, and trained with all samples except T1DGC. The number of individuals available for validation at each HLA locus is also shown. *Method does not support this locus.

| HLA locus | No. validation individuals | HIBAG | MAGprediction | HLA*IMP:02 |  | HLA*IMP:03 |  |
| --- | --- | --- | --- | --- | --- | --- | --- |
|  |  | Pre-trained | Pre-trained | CV | Excl. T1DGC | CV | Excl. T1DGC |
| *A* | 5,192 | 95.2 | 41.2 | 97.4 | 95.5 | 97.5 | 96.6 |
| *B* | 5,192 | 91.5 | 46.6 | 93.3 | 91.5 | 95.1 | 93.8 |
| *C* | 5,192 | 97.2 | 56.2 | 98.3 | 97.1 | 98.4 | 97.7 |
| *DPA1* | 5,192 | * | * | 99.7 | 98.3 | 99.8 | 98.8 |
| *DPB1* | 5,190 | 89.9 | 86.9 | 95.0 | 88.4 | 95.6 | 92.3 |
| *DQA1* | 5,192 | 98.4 | * | 98.8 | 98.2 | 99.4 | 98.8 |
| *DQB1* | 5,192 | 97.1 | 67.2 | 97.5 | 95.3 | 98.4 | 95.2 |
| *DRB1* | 5,192 | 88.8 | 37.6 | 93.6 | 90.1 | 95.3 | 93.7 |

The publicly available pre-trained versions of HLA*IMP:02 and HLA*IMP:03 could not be assessed in this manner since they include all of the T1DGC data in their reference panels. Instead we assessed two versions of these two methods: (i) five-fold CV; and (ii) all T1DGC data omitted from the reference panel. Since version (i) makes use of some T1DGC data to train each fold of CV, it may have some advantage due to ascertainment bias and not reflect performance in other independent European validation sets. This is not a concern for version (ii), however it may underestimate the performance of HLA*IMP:02 and HLA*IMP:03 relative to other methods in other independent validation sets, since the actual pre-trained versions have a larger reference panel. Thus these two versions could be considered as approximate lower and upper bounds on the relative performance of HLA*IMP:02 and HLA*IMP:03 to the other methods.

The CV versions of HLA*IMP:02 and HLA*IMP:03 were the best performing methods at each locus (with HLA*IMP:02 equal second with another method at some loci), and with HLA*IMP:03 better or equal to HLA*IMP:02 at each locus. HLA*IMP:03, with T1DGC data excluded from training, was better than pre-trained HIBAG at all loci except *HLA-DQB1*, and better than or equal to HLA*IMP:02, with T1DGC data excluded from training, at all loci. Pre-trained HIBAG outperformed HLA*IMP:02, with T1DGC data excluded from training, at *HLA-DQB1* and *HLA-DPB1*, but was substantially worse at *HLA-DQA1*.

Pre-trained MAGprediction was found to have extraordinarily poor accuracy. This seems to be due to different treatment of ambiguous alleles. For example, the alleles *A*02:01*, *A*03:01* and *A*24:02* (all ambiguous G alleles representing 25.8%, 11.8% and 11.6% of alleles in the T1DGC dataset, respectively) were most frequently imputed as *A*02:92*, *A*03:91* and *A*24:91*, respectively, by MAGprediction. (Each of these imputed alleles was not present in the merged dataset.) We did not attempt to investigate this further.

### HLA*IMP:03 in practical settings

The accuracy of HLA*IMP:03 using imputed SNPs (as described in the Material and Methods section) and imputed SNPs subset to those typed on common SNP arrays is provided in Table S6. The number of SNPs used for each SNP array is shown in Table S7. Using the extra imputed SNPs has no effect on accuracy. For most SNP arrays there is no loss of accuracy when using only SNPs typed on the array, and where there is loss of accuracy this is clearly explained by insufficient typing of SNPs across the xMHC (e.g. Affymetrix GeneChip Human Mapping 10K 2.0 and 100K Set, Illumina Human Omni1S-8 v1.0).

### Effect of call threshold on accuracy

Accuracy is plotted against call rate for each HLA locus in Figure S5. The accuracy at a call rate of 1 is the unthresholded accuracy (already presented above in Table 2 and Figure 1, but in this case aggregated over all populations). The plots demonstrate the trade-off between the accuracy of called imputations and call rate. The optimal call threshold depends on the intended use of the imputations (i.e. specification of a loss function) and is considered below for use in a statistical test of genetic association.

### Model calibration

The calibration of each method is shown in Figure S6. Well-calibrated predictions should have plotted points in the calibration plots that lie close to the diagonal line, although they may deviate by chance. This is taken into account by the credible interval, which for a well-calibrated model we expect to include the diagonal line at a rate of 95%. HLA*IMP:02 and SNP2HLA are the best-calibrated methods over the full range of posteriors, although HLA*IMP:02 tends to overestimate accuracy for higher posteriors. Both HIBAG and HLA*IMP:03 posteriors tend to underestimate accuracy, which in the case of HIBAG may be due to posteriors being provided per genotype rather than per allele (which is the basis on which accuracy was calculated). The distribution of posterior probabilities is shown in Figure S7. All methods have very high median posteriors, which is to be expected given that the methods are highly accurate and reasonably well-calibrated.

### Sample size ratio

We investigated the performance of HLA imputation when used in a test of genetic association for each allele (see Material and Methods) by considering the sample size ratio for each allele in the CV analysis with the multi-population reference panel, calculated using only the European individuals. We considered three versions of the sample size ratio based on: (i) unthresholded MAP imputations (call threshold *T* = 0), (ii) MAP imputations with call threshold that maximizes sample size ratio (optimal call threshold) set separately for each allele, and (iii) imputed dosage.

An illustrative example of how sample size ratio is affected by call threshold for a single allele (the most common allele of *HLA-B* in European individuals, *HLA-B*08:01*, allele frequency (AF) 13%) is provided in Figure S8. For this allele we observe a modest improvement in accuracy (sensitivity and specificity) with increasing call threshold that is not sufficient to compensate for the decreasing call rate, such that the optimal call threshold is 0.

The relationship between sample size ratio and call threshold is shown for all alleles of *HLA-B*, stratified by AF, in Figure S9. For common alleles (AF > 5%) the relationship between sample size ratio and call threshold is consistent and the optimal threshold is always very close to 0. For alleles of moderate frequency (1% < AF ≤ 5%) the relationship between sample size ratio and call threshold is smooth; the optimal threshold is variable across alleles but close to 0 for many alleles. For low frequency alleles (AF ≤ 1%) the relationship between sample size ratio and call threshold is often noisy, which is to be expected for the rarest alleles. The relationships were consistent across all imputation methods. The same relationships were also found for other HLA loci (data not shown).

The two versions of sample size ratio based on MAP imputations, unthresholded and with optimal call threshold, are compared for each imputation method and all HLA genes and alleles, stratified by AF, in Figure S10. For most common alleles (AF > 1%) using the optimal call threshold does not improve the sample size ratio. For low frequency alleles (AF ≤ 1%) the optimal call threshold does often improve sample size ratio, however the low AF means that power usually remains very low for these alleles, so there is little benefit in using the optimal call threshold (see illustrative example of power estimates below and Figure 3B).

**Figure 3.**
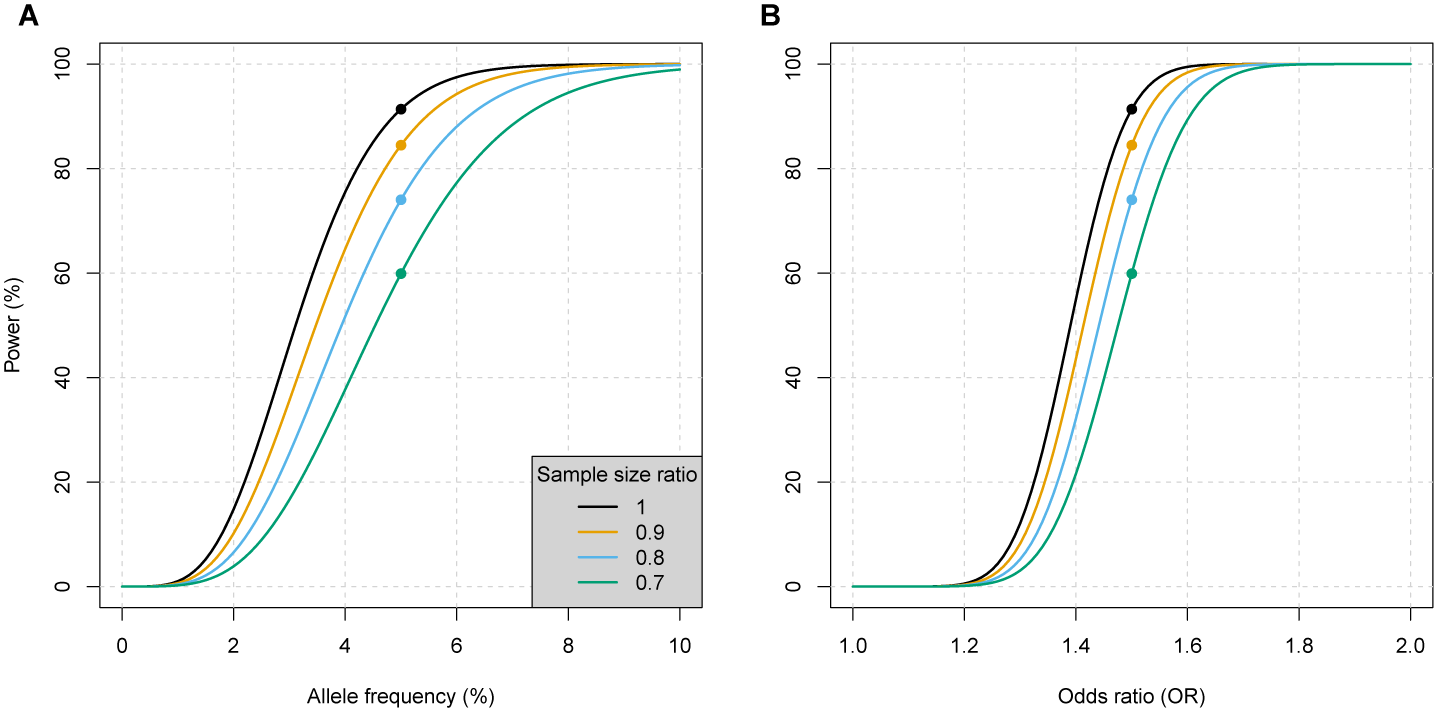
Impact of the sample size ratio on the power to detect an association. (**A**) Estimates of power for a standard association test (using the formula from Ref.^49^) conducted at a hypothetical allele for various assumed values of the sample size ratio and allele frequency. Further assumptions include: odds ratio (OR) of 1.5, a p-value threshold of 5 × 10^−8^, 5000 cases and 5000 controls. (**B**) Standard power curves for the same scenario as in (**A**) but this time varying the OR but keeping the allele frequency fixed at 5%. The points in each plot show the scenarios where the assumed parameter values are identical (OR of 1.5 and allele frequency 5%).

Sample size ratio based on dosages and MAP imputations with optimal threshold are similarly compared in Figure S11. Our observations are very similar; differences in sample size ratio are mainly observed for low frequency alleles, for which power will usually be low for both approaches. When there is a difference in sample size ratio at low frequency alleles the MAP imputations with optimal threshold are usually better. We note there are a number of very rare alleles present in the reference panel that are never imputed. The MAP sample size ratio has been set to zero for these alleles, reflecting the fact that power for detecting a genetic association is effectively zero. The dosage-based sample size ratio for these alleles is often non-zero.

Sample size ratios from both unthresholded MAP imputations and dosages, for alleles from all 11 imputed HLA loci with AF ≥ 1%, are compared pairwise between imputation methods in Figure S12. HIBAG and HLA*IMP:03 achieve best sample size ratios and are approximately equivalent in performance, both outperforming the other two methods. HLA*IMP:02 is the next best method, outperforming SNP2HLA.

Sample size ratio from dosages is plotted against AF, at all HLA loci in Figure S13. Sample size ratio is variable across AFs, although low sample size ratios occur more often for low frequency alleles. (The plots do not show alleles with AF < 1% for which sample size ratios can be very low.) The relative performance of imputation methods is as described above and consistent across alleles at a given HLA locus, although the magnitude of differences between methods is variable across loci.

To illustrate the impact of imputation directly on power we estimated power for a standard association test (using the formula from Ref.^49^) conducted at a hypothetical allele for a range of values of the sample size ratio (Figure 3A). We assumed an odds ratio of 1.5, a p-value threshold of 5 × 10^8^, 5000 cases and 5000 controls, and varied AF. Standard power curves for the same scenario but specifically for an AF of 5% are shown in Figure 3B. In this example we see how differences in sample size ratio, of the range observed in this study, translate into meaningful differences in power.

## Discussion

We have shown that the aggregation of the largest-to-date HLA reference panel leads to much higher imputation accuracy when used with leading methods. This ensures high throughput and accurate HLA typing, and consequently increased statistical power for association studies with HLA alleles. Our analysis has allowed comparison of the leading HLA imputation methods on the basis of: (i) accuracy achieved in Europeans with the currently available pre-trained methods; and (ii) accuracy when using each method with an identical reference panel. In the validation analyses we found consistent relative performance of the methods, assessed by accuracy at two-field resolution: HLA*IMP:03 is the best performing, closely followed by HIBAG, followed by HLA*IMP:02, while SNP2HLA had significantly worse performance. MAGprediction gave unsatisfactory performance, both in its inability to be run for a large reference panel and the low accuracy of the pre-trained method due to problematic allele codings.

Use of a multi-population reference panel has a positive or neutral effect on imputation accuracy of Europeans (depending on the method and HLA locus), however we did not determine whether imputation of non-Europeans is improved by inclusion of European reference samples. A multi-population panel is convenient as it does not require ascertainment of the ethnicity of study individuals and the use of different population-specific models.

Use of HLA*IMP:03 with increased SNP density from SNPs imputed in the reference panel did not have an effect on accuracy. This suggests that the use of overlapping SNPs when merging reference panels has not decreased SNP density yet to a level where imputation accuracy is degraded. We also found that HLA*IMP:03 is robust to the use of only those SNPs typed on common SNP arrays.

Our analysis with the sample size ratio showed that use of imputed dosages gives similar power in tests of genetic association to that obtained with the MAP imputed genotype, at least for alleles that are not too rare (frequency greater than 1%). In the context of SNP imputation, it has been shown both empirically^46^ and theoretically^47^ that the allelic dosage results in superior power, although in Ref.^46^ it was found that for high accuracy imputations (*r*^2^ > 0.9) there is little loss of power using the MAP imputations, and the gain from use of dosages is greatest at intermediate posteriors. In our context, the vast majority of imputations are done with high accuracy, which would explain why we saw that both approaches led to similar power. We recommend that if MAP imputations are used then no call threshold should be imposed, as generally the improvement in accuracy does not compensate for the reduction in call rate.

Imputation was computationally fastest for HIBAG and HLA*IMP:03 (a single analysis, e.g. single fold in CV for one HLA locus, took several minutes on a high performance cluster). Imputation for HLA*IMP:02 and SNP2HLA took longer (up to several hours), although for HLA*IMP:02 and HLA*IMP:03 imputations are performed on a web server, so there are no computational requirements for the user. Imputation with SNP2HLA was most burdensome due to severe memory requirements (more than 250 GB RAM in some cases), making its use with a reference panel of the size presented here impractical for most users. Moreover SNP2HLA is now provided with only a Pan-Asian reference panel, so imputation in other populations requires the user to obtain their own reference panel. (SNP2HLA is the only method that requires the reference panel in order to perform imputations; other methods allow use of only a pre-trained statistical model.) With the exception of SNP2HLA, training of the methods takes longer than imputation, taking several days for HIBAG and HLA*IMP:02 for a single analysis, but less than an hour for HLA*IMP:03. We note that HLA*IMP:02 and HLA*IMP:03 have not been made available for users to train with their own reference panel.

The difference in accuracy obtained for European and non-Europeans demonstrates the importance of large numbers of reference samples that match the target population. This is also shown by comparison with previous studies. The accuracy of HIBAG was assessed using approximately 3000 Japanese individuals, split into equal-sized training and validation sets,^50^ with accuracy in the range of 95.7% to 98.9% for *HLA-A*, -*B*, -*DRB1*, -*DQB1* and -*DPB1*. In the CV analysis, accuracy for the 80 JPT individuals in the 1000 Genomes dataset at these loci was in the range 87.5% to 93.6% for HIBAG and 89.4% to 94.9% for HLA*IMP:03 (the best performing method, see Table S4). This demonstrates the gain in accuracy that can be achieved with a larger number of reference samples that match the study panel.

Imputations using HLA*IMP, HLA*IMP:02 and SNP2HLA for *HLA-DRB1* were assessed in 161 Finnish individuals, with all methods achieving accuracy no greater than 25% at two-fields,^51^ however the pre-trained models used included only three Finnish individuals. In contrast, in the five-fold CV we found that accuracy with HLA*IMP:03 was 99.5% at *HLA-DRB1* for the 92 Finnish individuals in the 1000 Genomes dataset, and at least 96.2% at the other four typed HLA loci (*HLA-A*, -*B*, -*C* and -*DQB1*, see Table S4).

The mapping of HLA alleles to amino acid sequences^19^ allows association testing of individual amino acid sites. We note that this functionality is implemented in the HLA*IMP:03 web server. The HLA*IMP:03 web server also includes a tool for the calculation of allele-specific power based on sample size ratio from the CV analysis, with user specified values of OR, p-value threshold and sample size.

Finally, we note that more sophisticated treatment of ambiguous HLA alleles is likely to lead to further improvement in HLA imputation accuracy. The extent of allelic ambiguity can vary between independently typed samples (which was the case here for *HLA-DQA1*), and thus we advise that caution should be taken when making inferences based on imputation of potentially ambiguous HLA alleles.

## Appendix A: Sample size ratio for thresholded calls

Here we derive the *effective sample size* and *sample size ratio* of an association test that uses (imperfectly) imputed genotypes, and relate them to standard imputation accuracy measures. Where the goal of imputation is to conduct an association study, these are the most relevant measures of the effectiveness of imputation, rather than raw imputation accuracy.

As explained in the Material and Methods, a test of association considering a single allele with a presence/absence coding is equivalent to a biallelic marker. We can therefore use results derived for such markers. We follow closely the derivation for SNPs by Ref.;^52^ these describe a model of two SNPs in linkage disequilibrium, but the results apply equally well here for comparing two sets of biallelic types that are meant to be correlated. Specifically, let *A* represent the true type and *B* the imputed type at the locus of interest. The results of Ref.^52^ then imply that the effective sample size for a standard trend test is *N_A_* = *N_B_r*^2^, where *N_B_* is the actual study sample size and *r*^2^ is Pearson’s correlation coefficient comparing the imputed and true calls. This assumes no missing calls in the sample. If we impose a call threshold, resulting in a call rate of *c*, then we only have *N_B_c* of the sample remaining. Applying the above result to this reduced sample gives the overall effective sample size as *N_B_cr*^2^.

Both of these two multiplicative factors depend on the call threshold. They represent different and (typically) opposing effects. The call rate, *c*, will decrease as the threshold becomes more stringent, due to the exclusion of (hopefully) the less certain imputations. Conversely, the correlation, *r*^2^, should increase if the remaining imputations are indeed more accurate. The above formula shows us how to combine these two factors in order to assess their joint effect on power.

The correlation is a natural parameter when considering two SNPs in LD, but is not so natural in the context of imputation. We can re-express it in more standard quantities. Adopting the notation from Ref.^52^ and letting the allele of interest be coded as 1 and the (pooled) remaining alleles as 0, the true allele frequency (amongst the genotypes with non-null calls) is *f_A_* = Pr(*A* = 1), the allele frequency amongst the imputation calls is *f_B_* = Pr(*B* = 1), and we also have *q*_0_ = Pr(*A* = 1 | *B* = 0) and *q*_1_ = Pr(*A* = 1 | *B* = 1). The correlation is then a function of these, 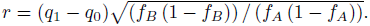. Rather than *q*_0_ and *q*_1_, here it is more natural to use a parameterisation in terms of the *sensitivity*, *s* = Pr(*B* = 1 | *A* = 1), and the *specificity, t* = Pr(*B* = 0 | *A* = 0). It can be shown that *q*_1_ = *sf_A_*/*f_B_*, *q*_0_ = (1 – s)*f_A_*/(1 – *f_B_*) and *f_B_* = (1 – *t*)(1 – *f_A_*) + *sf_A_*, which after some manipulation gives,

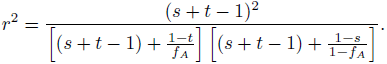

Therefore, the overall sample size ratio is,

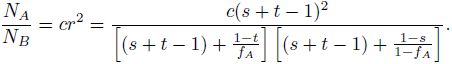

In principle, we could use the above expression to calculate the sample size ratio for a given application of HLA imputation, by providing assumed values or estimates of each of the quantities (allele frequency, call rate, sensitivity and specificity). Here we have taken the simpler approach of simply calculating the correlation between imputed and true types from the validation analyses. We did this separately for each of the four population groups, to provide estimates relevant for imputation within each of these populations.

## Conflict of Interest

A.D., P.D., G.M. and S.L. are partners in Peptide Groove LLP. Peptide Groove has licensed HLA typing technology to Affymetrix Ltd.

## Supplemental Data

Supplemental Data include 13 figures and 7 tables.

## Acknowledgments

We wish to thank the Type 1 Diabetes Genetics Consortium for providing the T1DGC dataset, David Morris for providing a dataset of African-American individuals, and Ingrid Kockum for providing a dataset of Swedish individuals.

This work was supported by the Australian National Health and Medical Research Council (NHMRC), Career Development Fellowship ID 1053756 (S.L.); and by the Victorian Life Sciences Computation Initiative (VLSCI) grant number VR0240 on its Peak Computing Facility at the University of Melbourne, an initiative of the Victorian Government, Australia (S.L.). Research at the Murdoch Childrens Research Institute was supported by the Victorian Government’s Operational Infrastructure Support Program.

## Web Resources

1000 Genomes Project SNP genotypes, ftp://ftp.1000genomes.ebi.ac.uk/vol1/ftp/technical/working/20110117_bi_omni_intensities/Omni25_genotypes.b36.vcf.gz

1958 Birth Cohort, http://www.b58cgene.sgul.ac.uk/

Affymetyrix, http://www.affymetrix.com/

Genome Reference Consortium GRCh37, https://www.ncbi.nlm.nih.gov/projects/genome/assembly/grc/human/

HLA*IMP:03, http://imp.mcri.edu.au/

HLA Nomenclature for reporting of ambiguous allele typings

G Codes, http://hla.alleles.org/alleles/g_groups.html

P Codes, http://hla.alleles.org/alleles/p_groups.html

Illumina, http://www.illumina.com/

LiftOver software, https://genome.ucsc.edu/cgi-bin/hgLiftOver

Michigan Imputation Server, https://imputationserver.sph.umich.edu

OMIM, http://www.omim.org/

PLINK software, http://pngu.mgh.harvard.edu/purcell/plink/

R software, http://www.r-project.org/

Pan-Asian HLA reference panel, https://www.broadinstitute.org/mpg/snp2hla/data/SNP2HLA_package_v1.0.3.tar.gz

## References

[1] Erlich, H., Opelz, G., and Hansen, J. (2001). HLA DNA Typing and Transplantation. Immunity 14, 347–356.

[2] International Multiple Sclerosis Genetics Consortium, Wellcome Trust Case Control Consortium 2, Sawcer, S., Hellenthal, G., Pirinen, M., Spencer, C. C. A., Patsopoulos, N. A., Moutsianas, L., Dilthey, A., Su, Z., et al. (2011). Genetic risk and a primary role for cell-mediated immune mechanisms in multiple sclerosis. Nature 476, 214–219.

[3] Moutsianas, L., Jostins, L., Beecham, A. H., Dilthey, A. T., Xifara, D. K., Ban, M., Shah, T. S., Patsopoulos, N. A., Alfredsson, L., Anderson, C. A., et al. (2015). Class II HLA interactions modulate genetic risk for multiple sclerosis. Nature Genetics 47, 1107–1113.

[4] Australo-Anglo-American Spondyloarthritis Consortium (TASC), Wellcome Trust Case Control Consortium 2 (WTCCC2), Evans, D. M., Spencer, C. C. A., Pointon, J. J., Su, Z., Harvey, D., Kochan, G., Opperman, U., Dilthey, A., et al. (2011). Interaction between ERAP1 and HLA-B27 in ankylosing spondylitis implicates peptide handling in the mechanism for HLA-B27 in disease susceptibility. Nature Genetics 43, 761–767.

[5] Genetic Analysis of Psoriasis Consortium and Wellcome Trust Case Control Consortium 2. (2010). A genome-wide association study identifies new psoriasis susceptibility loci and an interaction between HLA-C and ERAP1. Nature Genetics 42, 985–990.

[6] Han, B., Diogo, D., Eyre, S., Kallberg, H., Zhernakova, A., Bowes, J., Padyukov, L., Okada, Y., González-Gay, M. A., Rantap-Dahlqvist, S., et al. (2014). Fine Mapping Seronegative and Seropositive Rheumatoid Arthritis to Shared and Distinct HLA Alleles by Adjusting for the Effects of Heterogeneity. The American Journal of Human Genetics 94, 522–532.

[7] Hirayasu, K., Ohashi, J., Kashiwase, K., Hananantachai, H., Naka, I., Ogawa, A., Takanashi, M., Satake, M., Nakajima, K., Parham, P., et al. (2012). Significant Association of KIR2DL3-HLA-C1 Combination with Cerebral Malaria and Implications for Co-evolution of KIR and HLA. PLoS Pathog 8, e1002565.

[8] The International HIV Controllers Study. (2010). The Major Genetic Determinants of HIV-1 Control Affect HLA Class I Peptide Presentation. Science 330, 1551–1557.

[9] Dunstan, S. J., Hue, N. T., Han, B., Li, Z., Tram, T. T. B., Sim, K. S., Parry, C. M., Chinh, N. T., Vinh, H., Lan, N. P. H., et al. (2014). Variation at HLA-DRB1 is associated with resistance to enteric fever. Nature Genetics 46, 1333–1336.

[10] Moutsianas, L., Enciso-Mora, V., Ma, Y. P., Leslie, S., Dilthey, A., Broderick, P., Sherborne, A., Cooke, R., Ashworth, A., Swerdlow, A. J., et al. (2011). Multiple Hodgkin lymphoma-associated loci within the HLA region at chromosome 6p21.3. Blood 118, 670–674.

[11] Gragert, L., Fingerson, S., Albrecht, M., Maiers, M., Kalaycio, M., and Hill, B. T. (2014). Fine-mapping of HLA associations with chronic lymphocytic leukemia in US populations. Blood 124, 2657–2665.

[12] Bharadwaj, M., Illing, P., Theodossis, A., Purcell, A. W., Rossjohn, J., and McCluskey, J. (2012). Drug Hypersensitivity and Human Leukocyte Antigens of the Major Histocompatibility Complex. Annu. Rev. Pharmacol. Toxicol. 52, 401–431.

[13] Leslie, S., Donnelly, P., and McVean, G. (2008). A Statistical Method for Predicting Classical HLA Alleles from SNP Data. The American Journal of Human Genetics 82, 48–56.

[14] Lessard, C. J., Li, H., Adrianto, I., Ice, J. A., Rasmussen, A., Grundahl, K. M., Kelly, J. A., Dozmorov, M. G., Miceli-Richard, C., Bowman, S., et al. (2013). Variants at multiple loci implicated in both innate and adaptive immune responses are associated with Sjgren’s syndrome. Nature Genetics 45, 1284–1292.

[15] Dilthey, A. T., Moutsianas, L., Leslie, S., and McVean, G. (2011). HLA*IMP–an integrated framework for imputing classical HLA alleles from SNP genotypes. Bioinformatics 27, 968–972.

[16] Fairfax, B. P., Makino, S., Radhakrishnan, J., Plant, K., Leslie, S., Dilthey, A., Ellis, P., Langford, C., Vannberg, F. O., and Knight, J. C. (2012). Genetics of gene expression in primary immune cells identifies cell type–specific master regulators and roles of HLA alleles. Nature Genetics 44, 502–510.

[17] Dilthey, A., Leslie, S., Moutsianas, L., Shen, J., Cox, C., Nelson, M. R., and McVean, G. (2013). Multi-Population Classical HLA Type Imputation. PLoS Comput Biol 9, e1002877.

[18] Zheng, X., Shen, J., Cox, C., Wakefield, J. C., Ehm, M. G., Nelson, M. R., and Weir, B. S. (2013). HIBAG' HLA genotype imputation with attribute bagging. The Pharmacogenomics Journal 14, 192–200.

[19] Jia, X., Han, B., Onengut-Gumuscu, S., Chen, W.-M., Concannon, P. J., Rich, S. S., Raychaudhuri, S., and de Bakker, P. I. (2013). Imputing Amino Acid Polymorphisms in Human Leukocyte Antigens. PLoS ONE 8, e64683.

[20] Li, S. S., Wang, H., Smith, A., Zhang, B., Zhang, X. C., Schoch, G., Geraghty, D., Hansen, J. A., and Zhao, L. P. (2010). Predicting multiallelic genes using unphased and flanking single nucleotide polymorphisms. Genetic Epidemiology 35, 85–92.

[21] Xie, M., Li, J., and Jiang, T. (2010). Accurate HLA type inference using a weighted similarity graph. BMC Bioinformatics 11, S10.

[22] Setty, M. N., Gusev, A., and Pe’er, I. (2010). HLA Type Inference via Haplotypes Identical by Descent. In Lecture Notes in Computer Science In Lecture Notes in Computer Science. (Springer Science+Business Media).

[23] Paunić, V., Steinbach, M., Madbouly, A., and Kumar, V. (2014). Amb-EM: a SNP-based prediction of HLA alleles using ambiguous HLA data. In Proceedings of the 5th ACM Conference on Bioinformatics Computational Biology, and Health Informatics-BCB’14. (Association for Computing Machinery (ACM)) pp. 104–113.

[24] Paunic, V., Steinbach, M., Kumar, V., and Maiers, M. (2012). Prediction of HLA Genes from SNP Data and HLA Haplotype Frequencies. In 2012 IEEE 12th International Conference on Data Mining Workshops. (Institute of Electrical & Electronics Engineers (IEEE) pp. 964–971.

[25] Raychaudhuri, S., Sandor, C., Stahl, E. A., Freudenberg, J., Lee, H.-S., Jia, X., Alfredsson, L., Padyukov, L., Klareskog, L., Worthington, J., et al. (2012). Five amino acids in three HLA proteins explain most of the association between MHC and seropositive rheumatoid arthritis. Nature Genetics 44, 291–296.

[26] Hughes, T., Coit, P., Adler, A., Yilmaz, V., Aksu, K., Dzgn, N., Keser, G., Cefle, A., Yazici, A., Ergen, A., et al. (2013). Identification of multiple independent susceptibility loci in the HLA region in Beçet’s disease. Nature Genetics 45, 319–324.

[27] Okada, Y., Kim, K., Han, B., Pillai, N. E., Ong, R. T.-H., Saw, W.-Y., Luo, M., Jiang, L., Yin, J., Bang, S.-Y., et al. (2014). Risk for ACPA-positive rheumatoid arthritis is driven by shared HLA amino acid polymorphisms in Asian and European populations. Human Molecular Genetics 23, 6916–6926.

[28] Wissemann, W. T., Hill-Burns, E. M., Zabetian, C. P., Factor, S. A., Patsopoulos, N., Hoglund, B., Holcomb, C., Donahue, R. J., Thomson, G., Erlich, H., et al. (2013). Association of Parkinson Disease with Structural and Regulatory Variants in the HLA Region. The American Journal of Human Genetics 93, 984–993.

[29] Okada, Y., Momozawa, Y., Ashikawa, K., Kanai, M., Matsuda, K., Kamatani, Y., Takahashi, A., and Kubo, M. (2015). Construction of a population-specific HLA imputation reference panel and its application to Graves’ disease risk in Japanese. Nature Genetics 47, 798–802.

[30] Goyette, P., Boucher, G., Mallon, D., Ellinghaus, E., Jostins, L., Huang, H., Ripke, S., Gusareva, E. S., Annese, V., Hauser, S. L., et al. (2015). High-density mapping of the MHC identifies a shared role for HLA-DRB1*01:03 in inflammatory bowel diseases and heterozygous advantage in ulcerative colitis. Nature Genetics 47, 172–179.

[31] Gabriel, C., Frst, D., Faé, I., Wenda, S., Zollikofer, C., Mytilineos, J., and Fischer, G. F. (2014). HLA typing by next-generation sequencing-getting closer to reality. Tissue Antigens 83, 65–75.

[32] Hosomichi, K., Shiina, T., Tajima, A., and Inoue, I. (2015). The impact of next-generation sequencing technologies on HLA research. Journal of Human Genetics 60, 665–673.

[33] Sudlow, C., Gallacher, J., Allen, N., Beral, V., Burton, P., Danesh, J., Downey, P., Elliott, P., Green, J., Landray, M., et al. (2015). UK Biobank: An Open Access Resource for Identifying the Causes of a Wide Range of Complex Diseases of Middle and Old Age. PLoS Med 12, e1001779.

[34] Vukcevic, D., Traherne, J. A., Næss, S., Ellinghaus, E., Kamatani, Y., Dilthey, A., Lathrop, M., Karlsen, T. H., Franke, A., Moffatt, M., et al. (2015). Imputation of KIR Types from SNP Variation Data. The American Journal of Human Genetics 97, 593–607.

[35] Levin, A. M., Adrianto, I., Datta, I., Iannuzzi, M. C., Trudeau, S., McKeigue, P., Montgomery, C. G., and Rybicki, B. A. (2014). Performance of HLA allele prediction methods in African Americans for class II genes HLA-DRB1-DQB1, and DPB1. BMC Genet 15, 72.

[36] The International HapMap Consortium. (2007). A second generation human haplotype map of over 3.1 million SNPs. Nature 449, 851–861.

[37] de Bakker, P. I. W., McVean, G., Sabeti, P. C., Miretti, M. M., Green, T., Marchini, J., Ke, X., Monsuur, A. J., Whittaker, P., Delgado, M., et al. (2006). A high-resolution HLA and SNP haplotype map for disease association studies in the extended human MHC. Nature Genetics 38, 1166–1172.

[38] Gourraud, P.-A., Khankhanian, P., Cereb, N., Yang, S. Y., Feolo, M., Maiers, M., Rioux, J. D., Hauser, S., and Oksenberg, J. (2014). HLA Diversity in the 1000 Genomes Dataset. PLoS ONE 9, e97282.

[39] The 1000 Genomes Project Consortium. (2012). An integrated map of genetic variation from 1,092 human genomes. Nature 491, 56–65.

[40] Mychaleckyj, J. C., Noble, J. A., Moonsamy, P. V., Carlson, J. A., Varney, M. D., Post, J., Helm-berg, W., Pierce, J. J., Bonella, P., Fear, A. L., et al. (2010). HLA genotyping in the international Type 1 Diabetes Genetics Consortium. Clinical Trials 7, S75–S87.

[41] Pillai, N. E., Okada, Y., Saw, W.-Y., Ong, R. T.-H., Wang, X., Tantoso, E., Xu, W., Peterson, T. A., Bielawny, T., Ali, M., et al. (2014). Predicting HLA alleles from high-resolution SNP data in three Southeast Asian populations. Human Molecular Genetics 23, 4443–4451.

[42] Howie, B., Fuchsberger, C., Stephens, M., Marchini, J., and Abecasis, G. R. (2012). Fast and accurate genotype imputation in genome-wide association studies through pre-phasing. Nature Genetics 44, 955–959.

[43] Delaneau, O., Zagury, J., and Marchini, J. (2013). Improved whole-chromosome phasing for disease and population genetic studies. Nat Methods 10, 5–6.

[44] Purcell, S., Neale, B., Todd-Brown, K., Thomas, L., Ferreira, M. A., Bender, D., Maller, J., Sklar, P., de Bakker, P. I., Daly, M. J., et al. (2007). PLINK: A Tool Set for Whole-Genome Association and Population-Based Linkage Analyses. The American Journal of Human Genetics 81, 559–575.

[45] Browning, B. L. and Browning, S. R. (2009). A Unified Approach to Genotype Imputation and Haplotype-Phase Inference for Large Data Sets of Trios and Unrelated Individuals. The American Journal of Human Genetics 84, 210–223.

[46] Zheng, J., Li, Y., Abecasis, G. R., and Scheet, P. (2011). A comparison of approaches to account for uncertainty in analysis of imputed genotypes. Genetic Epidemiology 35, 102–110.

[47] Liu, K., Luedtke, A., and Tintle, N. (2013). Optimal Methods for Using Posterior Probabilities in Association Testing. Human Heredity 75, 2–11.

[48] Howie, B. N., Donnelly, P., and Marchini, J. (2009). A Flexible and Accurate Genotype Imputation Method for the Next Generation of Genome-Wide Association Studies. PLoS Genetics 5, e1000529.

[49] Chapman, J. M., Cooper, J. D., Todd, J. A., and Clayton, D. G. (2003). Detecting Disease Associations due to Linkage Disequilibrium Using Haplotype Tags: A Class of Tests and the Determinants of Statistical Power. Human Heredity 56, 18–31.

[50] Khor, S.-S., Yang, W., Kawashima, M., Kamitsuji, S., Zheng, X., Nishida, N., Sawai, H., Toyoda, H., Miyagawa, T., Honda, M., et al. (2015). High-accuracy imputation for HLA class I and II genes based on high-resolution SNP data of population-specific references. Pharmacogenomics J 15, 530–537.

[51] Vlachopoulou, E., Lahtela, E., Wennerstrm, A., Havulinna, A. S., Salo, P., Perola, M., Salomaa, V., Nieminen, M. S., Sinisalo, J., and Lokki, M.-L. (2014). Evaluation of HLA-DRB1 imputation using a Finnish dataset. Tissue Antigens 83, 350–355.

[52] Vukcevic, D., Hechter, E., Spencer, C., and Donnelly, P. (2011). Disease model distortion in association studies. Genet Epidemiol 35, 278–90.

